# Optimization of conditions for conjugation of outer membrane vesicles of *Salmonella* Typhimurium with oral delivery systems using response surface method

**DOI:** 10.1101/2022.02.14.480461

**Authors:** Anisha Sultana, Rita Nath, Shantanu Tamuly, Yasmin Begum, Jayanta Sarmah Boruah, Arijit Shome, Devashish Chowdhury, Suraksha Subedi Deka, Dhruba Jyoti Kalita

## Abstract

**Background:** Outer membrane vesicles (OMVs) released from *Salmonella* Typhimurium can become the effective vaccine candidate. For efficient delivery through oral route, the OMVs are conjugated with some delivery systems. The current study was aimed at optimization of conditions required for conjugation of OMVs with nano- or microparticles for maximum entrapment of OMV in terms of protein concentration.

**Methods:** The OMVs of *Salmonella* Typhimurium were conjugated under three optimum conditions of pH, temperature and ratio (nanoparticles or microparticles: OMVs) predicted by response surface method. The efficiency of conjugation was determined by entrapment of OMVs in nano/microparticles.

**Result:** The pH and temperature were not influential conditions in case of conjugation of OMVs with chitosan nanoparticles (Ch-NP) and poly-lactide co-glycolide microparticles (PLG-MP) while they had influence in case of poly(anhydride) nanoparticles. In case of Ch-NP and PLG-MP, the optimum ratio for maximum entrapment was found to be 1:10 and 1:9 respectively. The optimum pH and temperature was found to be 7 and 24°C respectively for conjugation of poly(anhydride) nanoparticles. The optimized conditions did not alter the protein profile and immunogenic potential of conjugated vaccines.

## INTRODUCTION

*Salmonella*, a genus of flagellated rod-shaped, gram-negative, facultatively anaerobic bacteria, has been recognized as the food-borne pathogen of both humans and warm-blooded animals. The domestic poultry act as the largest reservoir for *Salmonella enterica* serotype Typhimurium. In humans and cattle, *S*. Typhimurium is responsible for diarrheal enterocolitis without systemic symptoms in majority of the cases. The Gram-negative bacteria shed outer membrane blebs, called native outer membrane vesicles (NOMVs) which are small, spherically bilayered with a size that ranges from 10-300 nm. The OMVs has been reported to be an efficient vaccine candidate (Alaniz et al., 2007). These OMVs do not require adjuvant if administered parenterally. However, if they are administered orally, they would require oral delivery systems. Nanoparticles and microparticles, due to their mucoadhesive properties, can play the role of delivery systems for the OMVs when administered as oral vaccine. In many formulations, the vaccine antigens are added during the formation of nanoparticles (He et al., 2000, 2002; Tamuly et al., 2014), however it is not feasible with all nanoparticles as many of them are made in harsh conditions that may reduce the immunopotential of entrapped antigen. The safest method appears to incorporation of antigens directly to preformed nanoparticles. For efficient formulation nanoparticle/microparticle based vaccine, it is of paramount importance to optimize the conditions for efficient conjugation of antigens with the delivery systems without compromising the immunopotential.

In this study, we optimized the conditions for conjugation of chitosan nanoparticles, poly-lactide co-glycolide microparticles and poly(anhydride) nanoparticles with OMVs of *Salmonella* Typhimurium using response surface method (RSM).

## MATERIALS AND METHODS

### Bacterial strain

The *Salmonella* Typhimurium MTCC-98 strain, was procured from Department of Animal Biotechnology, Assam Agricultural University, Khanapara, Guwahati, Assam.

### Isolation of OMVs

The OMVs were purified as per the method described by Solo et al. (2021). The protein concentration of the isolated OMVs was estimated using Lowry’s method (Lowry et al., 1951).

### Preparation of PLG microparticles

To 10.8 mg of PLG (50:50) dissolved in 60 μL dichloromethane, 6.4 μL of distilled water was added and sonicated for 20 seconds. The 1.296 ml of 2% polyvinyl alcohol was added and sonicated again for 5 minutes. After overnight stirring of the suspension, centrifugation was carried out at 10,000×g for 5 minutes followed by lyophilization.

### Preparation of poly (methyl vinyl ether-alt-maleic anhydride) nanoparticles

The poly(anhydride) nanoparticles were prepared following the method described by Camacho et al. (2011). The nanoparticles were lyophilized till further use.

### Chitosan nanoparticles

The chitosan nanoparticles were provided by Dr. Devasish Chowdhury, Institute of Advanced Study in Science and Technology (IASST), Guwahati, Assam.

### Characterization of nanoparticles and microparticles

The nanoparticles and microparticles were characterized with regards to zeta size and zeta potential by dynamic light scattering in IASST.

### Screening experiment

In the present study, the three conditions *viz*., pH, temperature and nanoparticles or microparticles:OMVs ratio (P:OMV) were taken to assess their influence in conjugation. The screening of these conditions were carried out by taking two levels of each of the conditions for the three adjuvants as depicted in table 1. The levels of different variables were coded using the formulae as given below:

Coded value= (Real value — central value)/ (0.5 × Range)
Central value= (Highest level + lowest level)/2
Range= Highest level + lowest level

**Table 1:** Full factorial experimental design for screening of the three factors.

| Sl. No. | Coded values |  |  | Real values |  |  |
| --- | --- | --- | --- | --- | --- | --- |
|  | pH | Temperature | Ratio | pH | Temperature (°C) | Ratio |
| 1 | -1 | -1 | -1 | 6.5 | 20 | 1:1 |
| 2 | +1 | -1 | -1 | 7.5 | 20 | 1:1 |
| 3 | -1 | +1 | -1 | 6.5 | 25 | 1:1 |
| 4 | +1 | +1 | -1 | 7.5 | 25 | 1:1 |
| 5 | -1 | -1 | +1 | 6.5 | 20 | 1:5 |
| 6 | +1 | -1 | +1 | 7.5 | 20 | 1:5 |
| 7 | -1 | +1 | +1 | 6.5 | 25 | 1:5 |
| 8 | +1 | +1 | +1 | 7.5 | 25 | 1:5 |

**Table 2:** Assessment of immunopotency of different OMV-nanoparticle conjugates.

| Groups of mice | Vaccines | Vaccine formulations |
| --- | --- | --- |
| Group I | poly(anhydride) nanoparticles<br>nanoparticle-OMV | Control condition |
| Group II | poly(anhydride) nanoparticles<br>nanoparticle-OMV | Predicted using RSM |
| Group III | PLG microparticles-OMV | Control conditions |
| Group IV | PLG microparticles-OMV | Predicted using RSM |
| Group V | Chitosan nanoparticle-OMV | Control conditions |
| Group VI | Chitosan nanoparticle-OMV | Predicted using RSM |
| Group VII | PBS | Unimmunized control |

### Conjugation of nanoparticles /microparticles with OMVs

The chitosan nanoparticles or PLG microparticles or poly(anhydride) nanoparticles were conjugated with the OMVs according to the conditions mentioned in the Table 1. The period of conjugation was for two hours. The suspension was centrifuged at 15,000×g for 10 minutes at 4 °C. The protein concentration of the supernatant was estimated in three replicates by Lowry’s method. The entrapment efficiency of OMVs in the nanoparticles/microparticles was calculated using the formulae mentioned below.

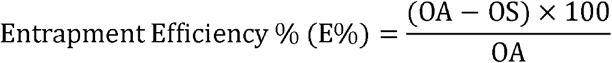

OA= amount of OMVs taken for conjugation

OS= amount of OMVs in the supernatant.

### Optimization of conditions by response surface methodology

The factors that were found to have significant influence on conjugation were taken further for optimization through response surface methodology (RSM). The 2^k^ design was used for determination of optimum condition (k=no. of factors). The entrapment efficiency nanoparticles/ microparticles was taken as response variable. For revalidation of optimized conditions, the experiments were set up in the predicted optimum conditions

### Immunopotential studies of vaccines in mice

The mice were immunized with the vaccine formulations prepared from the optimized conditions and the control condition of ratio 1:1, pH=7.0 and temperature=25 °C. The mice were immunized at the rate of 50 μg per mice. The blood samples were collected on 7 and 14 days post-primary immunization. The anti-OMV IgM response was measured using indirect ELISA using OMV as an antigen and the serum of control mice as the control serum. The ELISA was carried out as per the method described by Liu *et al*. (2016).

The protein profiling of the vaccine formulations was determined by sodium dodecyl sulfate polyacrylamide gel electrophoresis (SDS-PAGE) (Sambrook & Russels, 2001)

### Statistical analysis

The statistical data analysis was performed using statistical software R version 4.0.2 (R Core Team, 2020). To assess the significant influence of different factors on the entrapment efficiency, linear regression method was applied. The condition(s) with higher value of slope were considered as significantly influential. The central composite design was used when the number of influential conditions (factors) were more than 1 for RSM. The analysis was performed in rsm (Lenth, 2009) and pid packages (Dunn, 2018) of the R statistical software. For revalidation of the optimized conditions, the difference between the predicted and experimentally determined entrapment efficiency was assessed using one sample t-test. The difference in immune response in mice was analyzed by t-test. The p<0.05 was considered statistically significant.

## RESULTS AND DISCUSSION

In the current study, the influential factors determined for conjugation of nanoparticles/microparticles with OMVs were validated experimentally in terms of entrapment efficiency.

### Characterization of nanoparticles/ microparticles

The zeta size of PLG microparticles, chitosan nanoparticles and poly(anhydride) nanoparticles were found to be 3.6 ± 0.9 μm, 130.4 ± 16 nm and 247. 4 ± 14.3 nm, respectively and zeta potential of 3.38 mV, 12.5 mV and —52.6 mV respectively.

### Conjugation

Upon screening, the factors pH and temperature were found to have no significant influence in the entrapment efficiency of PLG microparticles but have significant influence in case of poly(anhydride) nanoparticles (Fig 4). As the pKa value of chitosan is ~6.5, its positive charge did not alter significantly in the selected pH range in our study (Abouelmagd et al., 2015). In case of PLG, the hydroxyl groups do not alter significantly in the pH range of 6.5 to 7.5 hence it did not influence the conjugation (Croll et al., 2004). The Fig 4 and 5 indicates that the chitosan nanoparticles/PLG microparticle to OMV ratio had significantly positive influence in entrapment efficiency within the range of 1:1-1:5. The Fig 6 indicates that the temperature had significantly positive influence and pH had significantly negative influence in entrapment efficiency within the ranges of 20-25°C and 6.5-7.5, respectively in case of poly(anhydride) nanoparticles. The alteration of pH and temperature greatly influence the surface properties of poly(anhydride) nanoparticles. We could not find any literature supporting this finding however it may be assumed that as the pK_2_ value of maleic acid component of poly(anhydride) nanoparticles is 6, hence it remains ionized at the range of pH 6.5 to 7.5 (Mohanty et al., 2013) and this ionization is probably influenced by the temperature. The ionized state of poly(anhydride) nanoparticles are important for their entrapment efficiency (Camacho et al., 2011).

**Fig 1:**
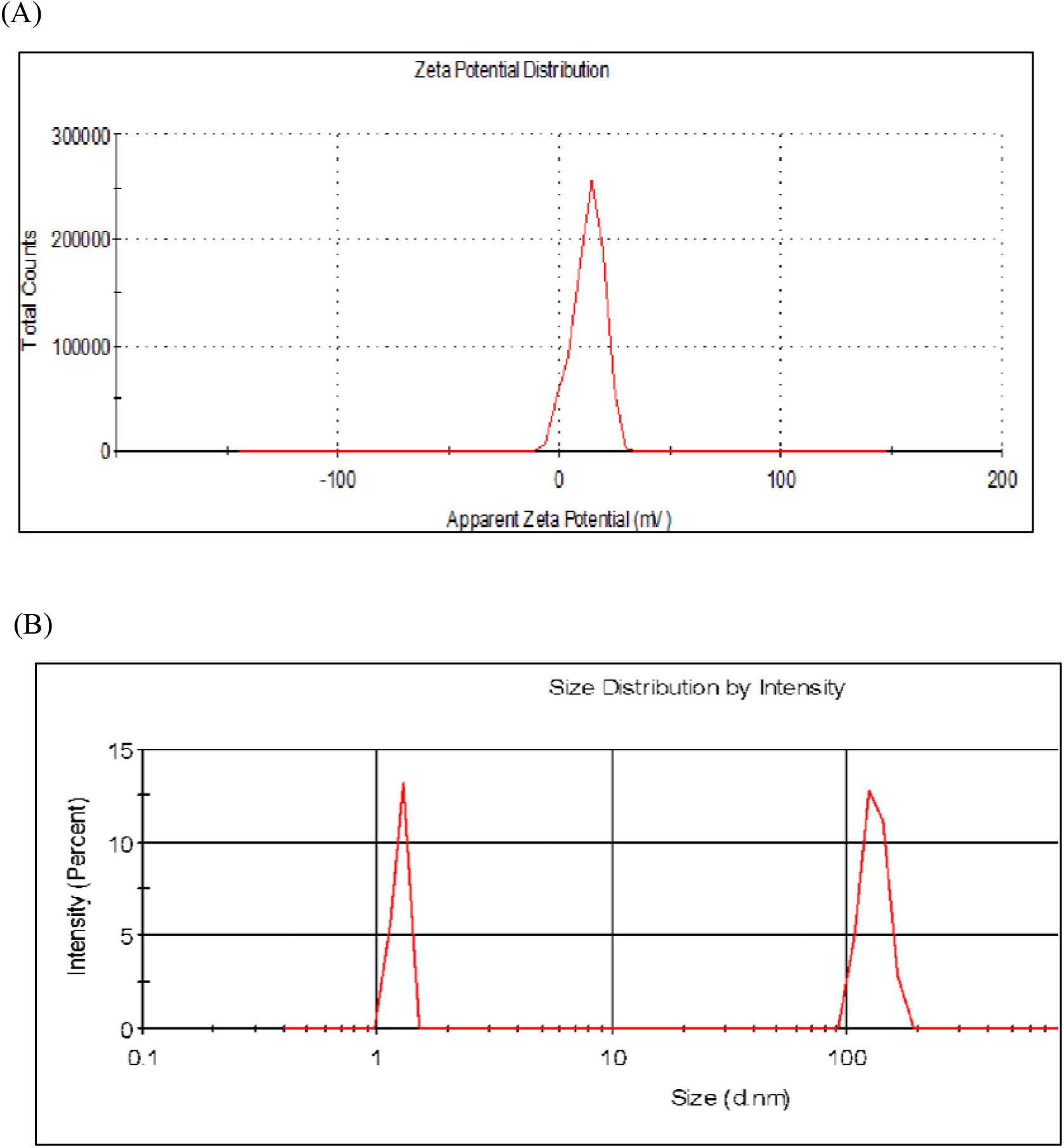
(A) Zeta Potential and (B) Zeta Size of chitosan nanoparticles.

**Fig 2:**
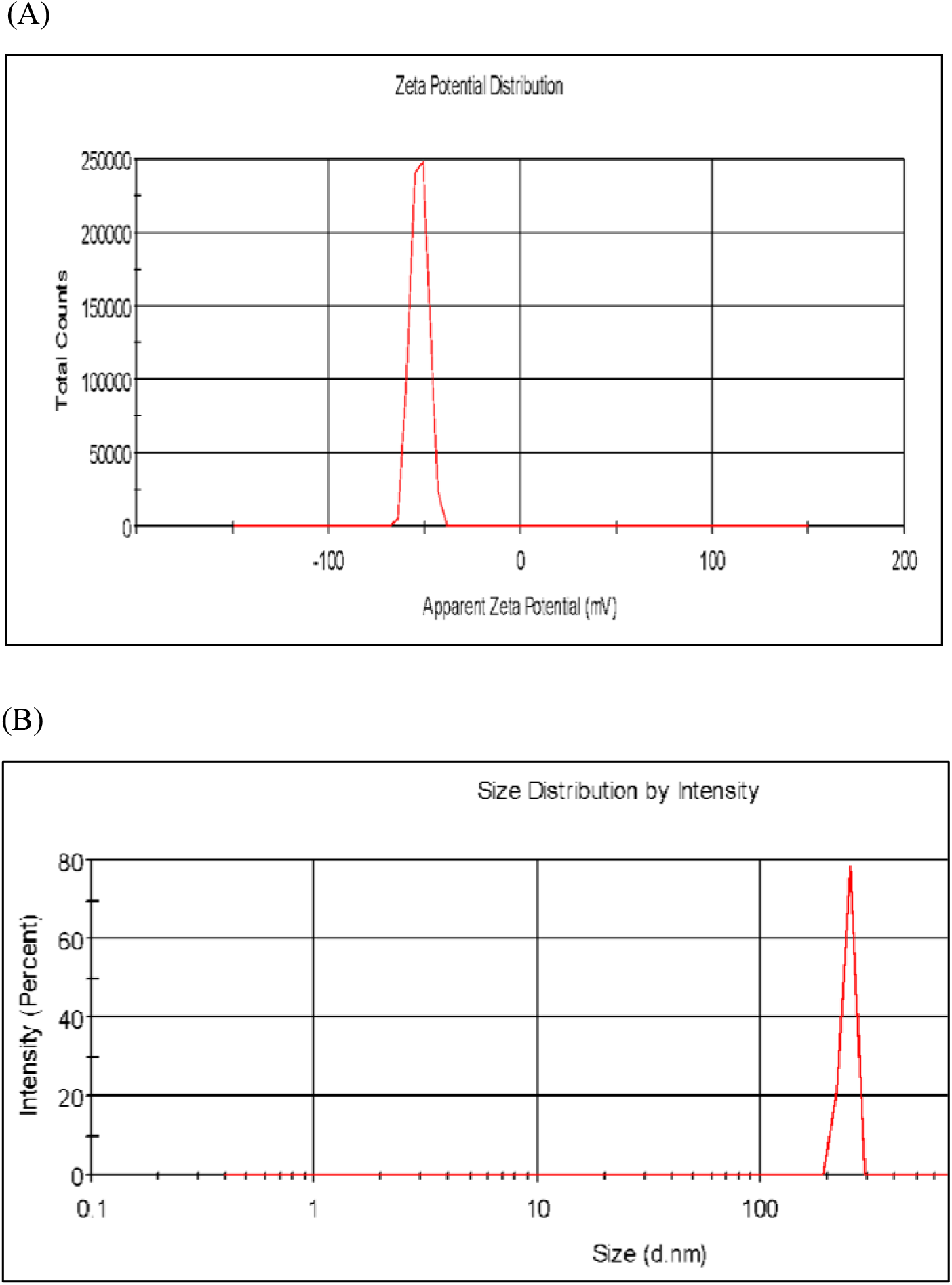
(A) Zeta Potential and (B) Zeta Size of poly(anhydride) nanoparticles.

**Fig 3:**
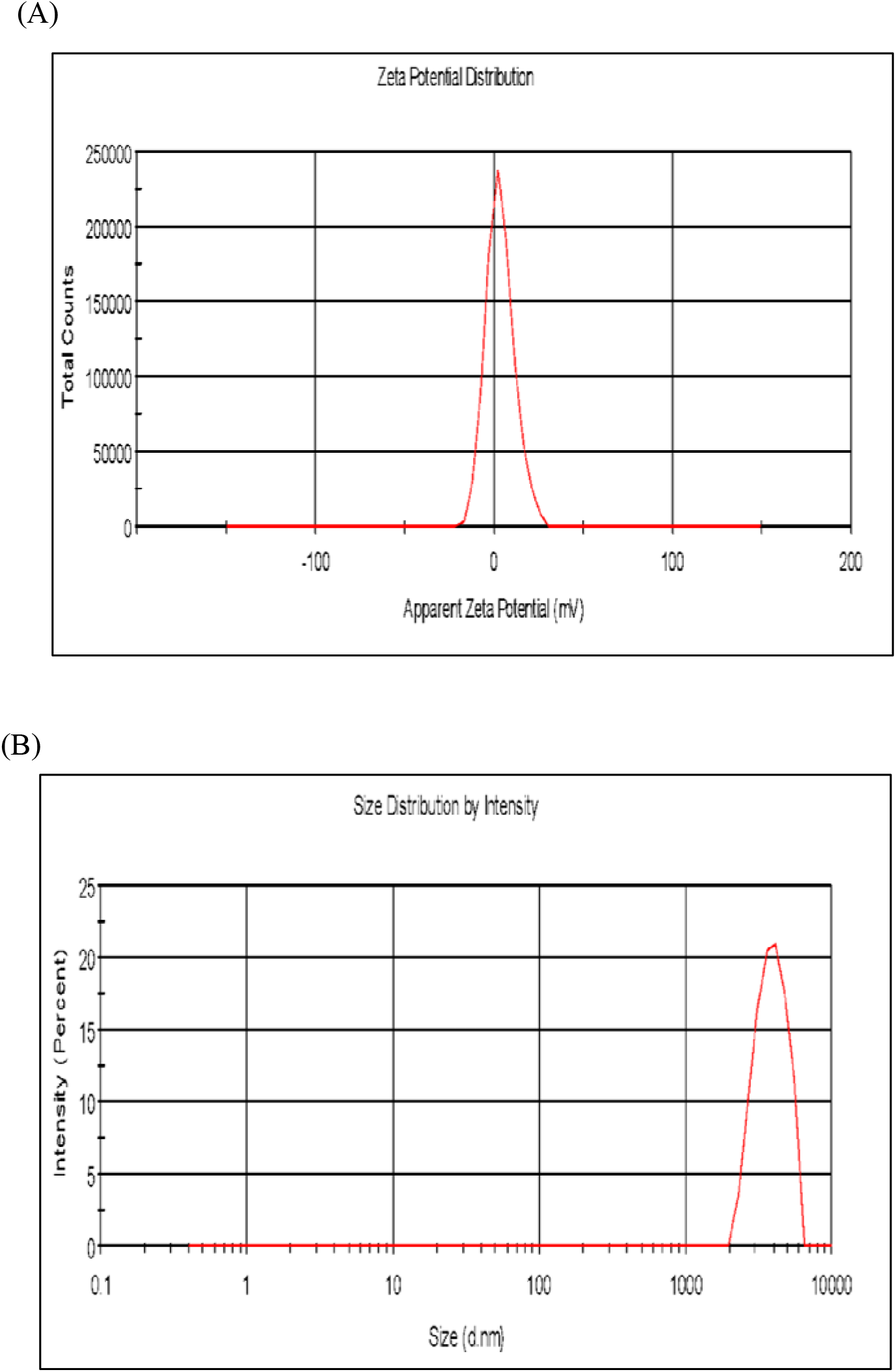
(A) Zeta Potential and (B) Zeta Size of PLG microparticles.

**Fig 4:**
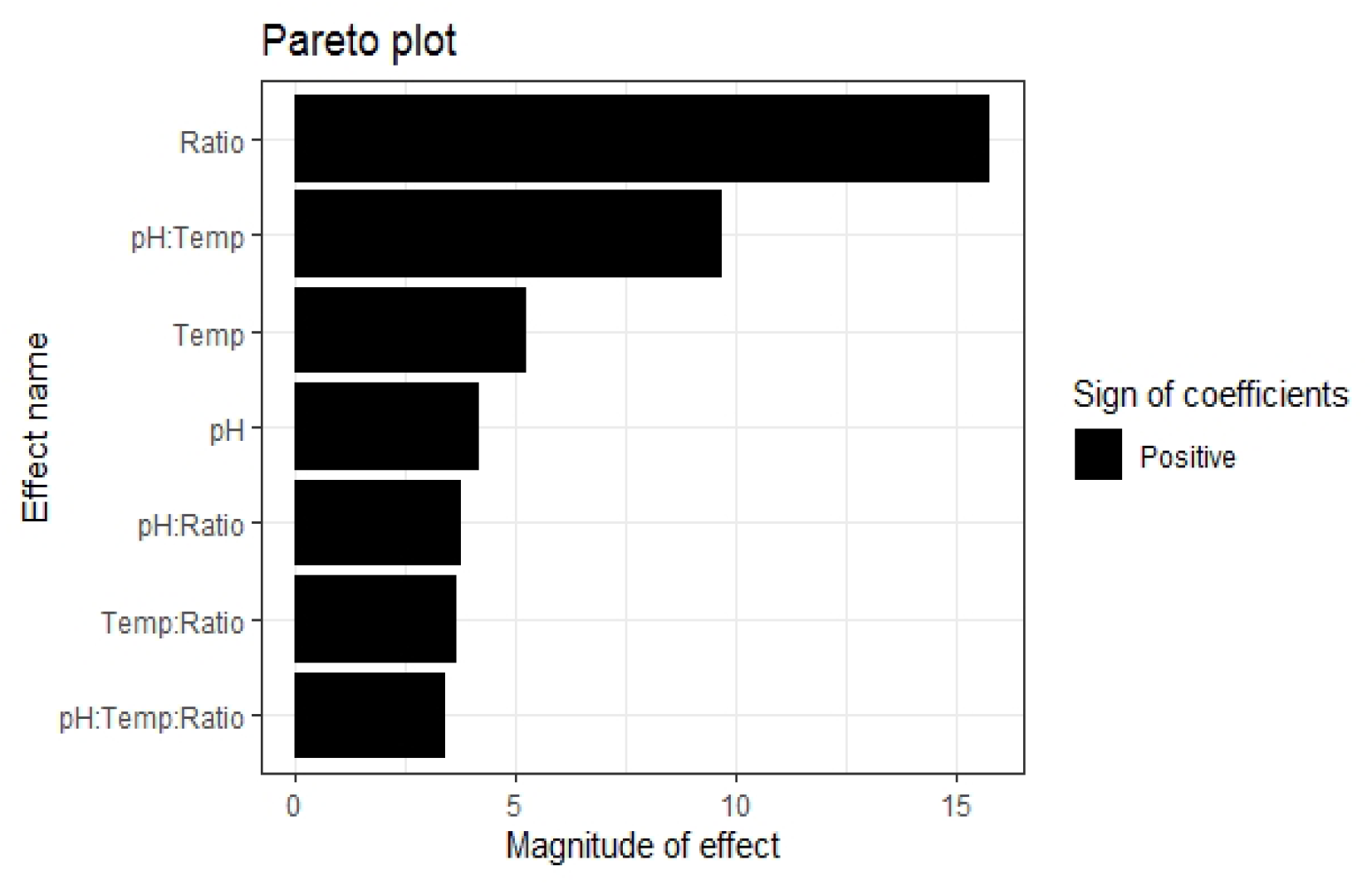
Pareto plot presenting the influence of different factors, *viz*. pH, chitosan nanoparticles: OMV ratio and temperature during conjugation.

**Fig 5:**
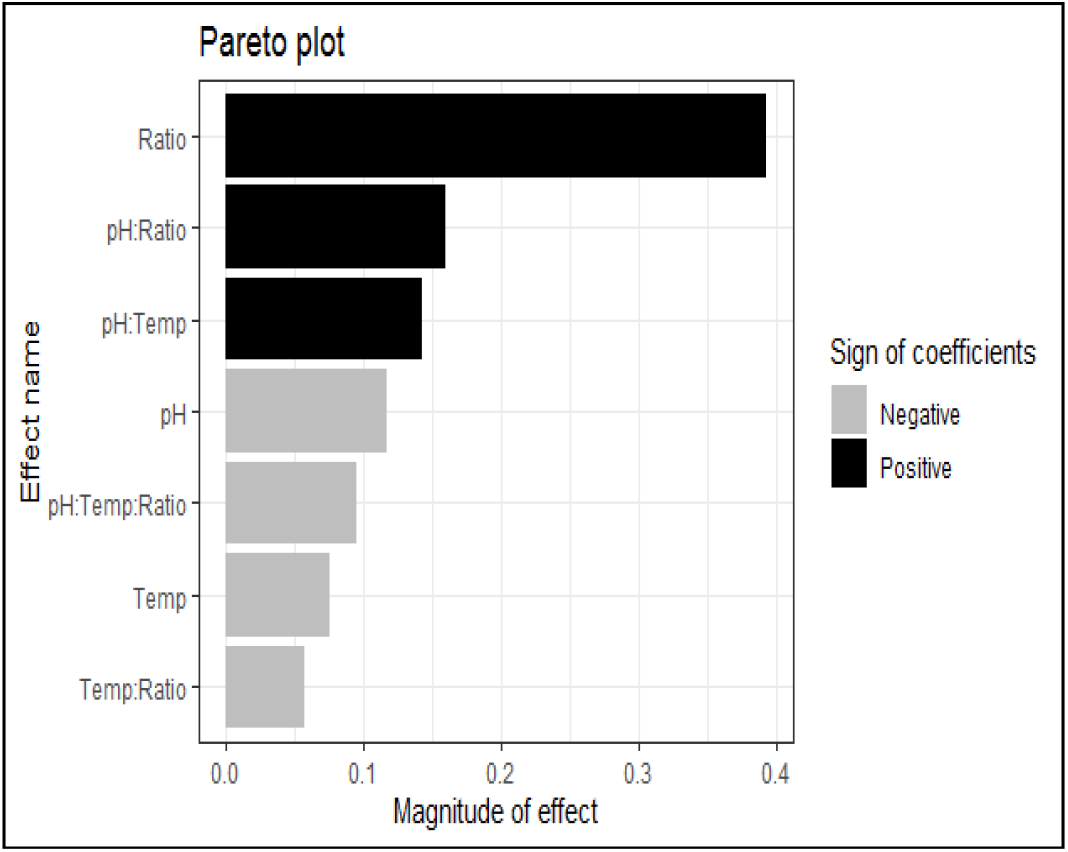
Pareto plot presenting the influence of different factors, *viz*. pH, PLG microparticles: OMV ratio and temperature during conjugation.

**Fig 6:**
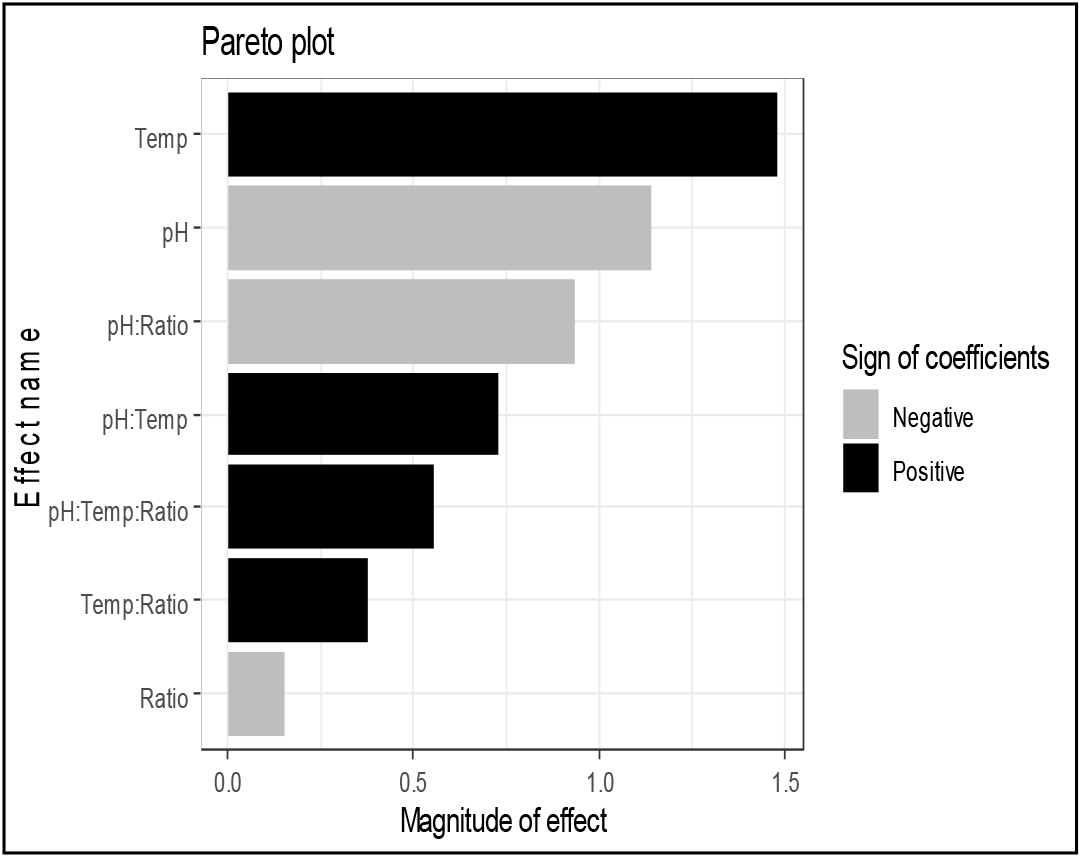
Pareto plot presenting the influence of different factors, *viz*. pH, poly(anhydride) nanoparticles: OMV ratio and temperature during conjugation.

### Conditions for determination of maximum OMV protein entrapment efficiency into nanoparticles/ microparticles

The P:OMV ratio ranging from 1:1 to 1:11 was taken for conjugation of OMV with PLG-MP and Ch-NP at constant temperature of 24°C and pH of 7.0. The pH and temperature ranging from 6.5 to 7.5 and 20 to 25°C respectively were taken for conjugation of OMV with poly(anhydride) nanoparticle at the constant P:OMV ratio of 1:1. The conjugation was carried out for 2 hours followed by centrifugation at 1000×g for 5 minutes. The entrapment efficiency of nanoparticles/microparticles was determined as mentioned earlier. The second order regression models with interaction terms were developed that was used for prediction of optimum levels of process variables for yielding maximum entrapment of OMV into nanoparticle/microparticles. The optimum ratio for Ch-NP—OMV and PLG-MP—OMV conjugation was predicted to be 1:10 (R^2^ =90%, p<0.05) and 1:9 (R^2^=91%, p<0.05) respectively as depicted in the Fig 7 and 8. On the other hand the optimum conditions for poly(anhydride) nanoparticle—OMV conjugation are pH=7.0 and temperature=24°C (R^2^=69%, p<0.05) as depicted in Fig 9. The models of conjugation can be summarized as below:

**Fig 7:**
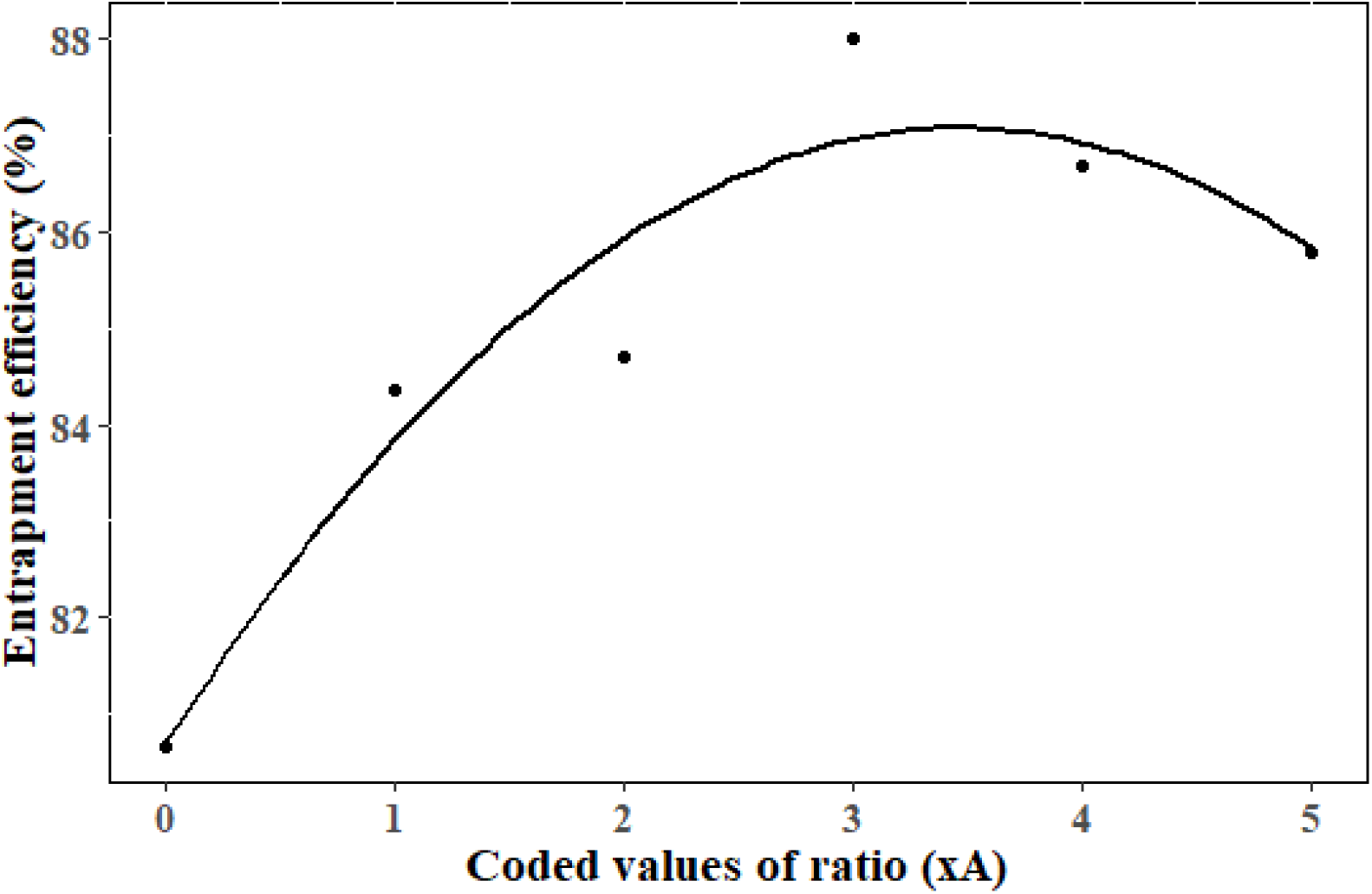
The response surface curve of entrapment efficiency of chitosan nanoparticles and OMV in different ratios.

**Fig 8:**
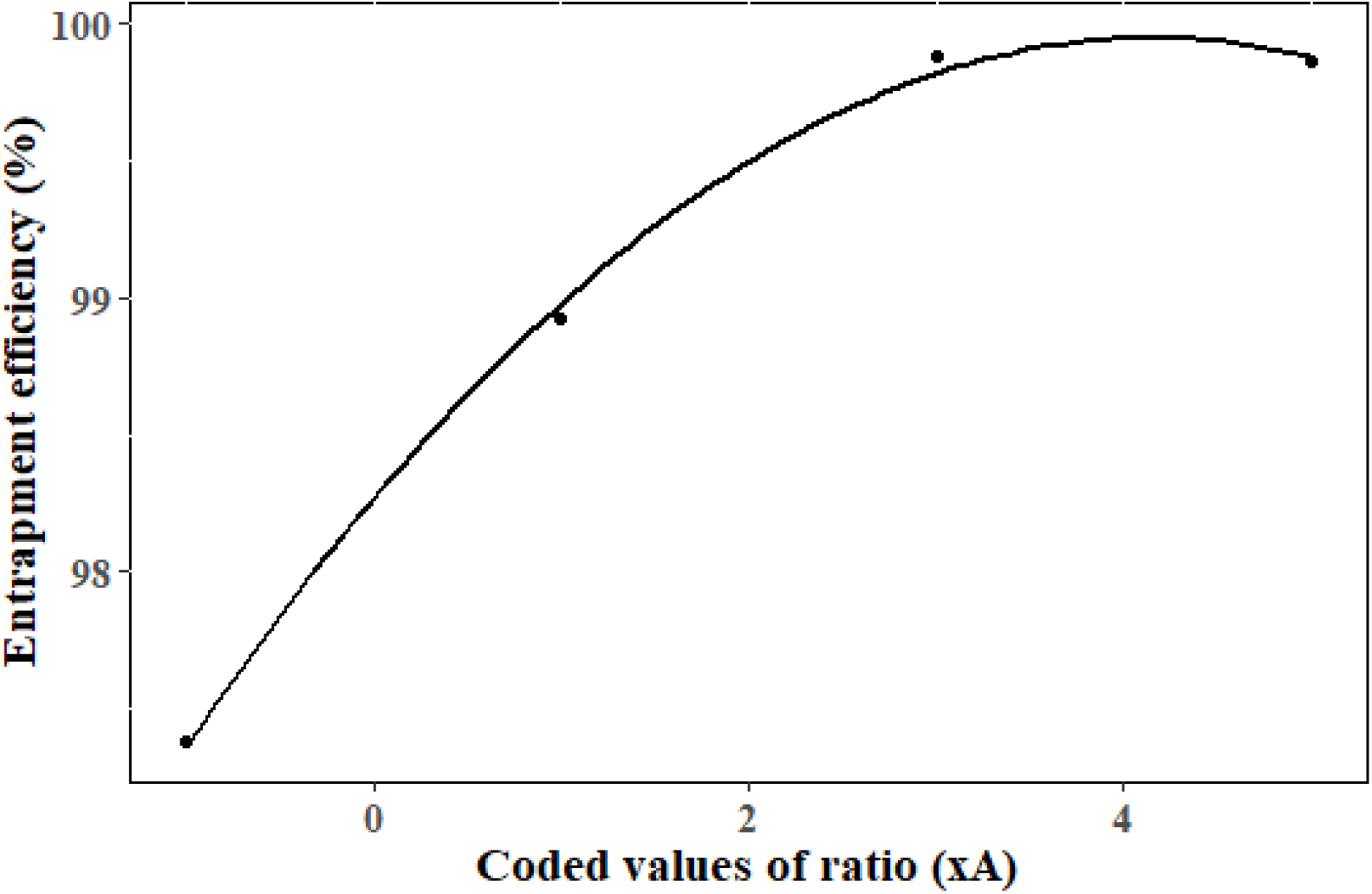
The response surface curve of entrapment efficiency of PLG microparticles and OMV in different ratios.

**Fig 9:**
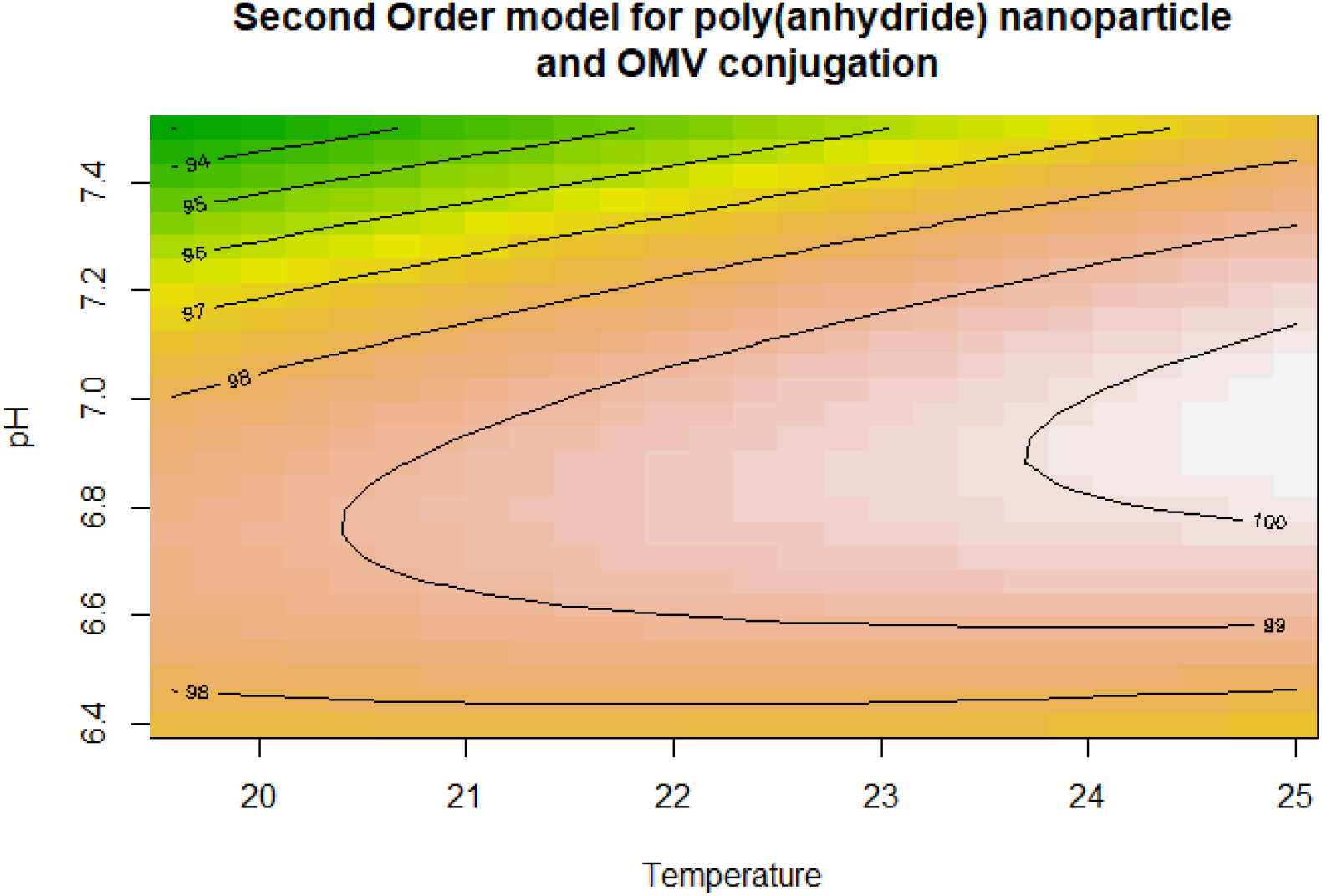
The contour plot of entrapment efficiency of poly(anhydride) nanoparticles and OMV in different pH and temperatures.

**Fig 10:**
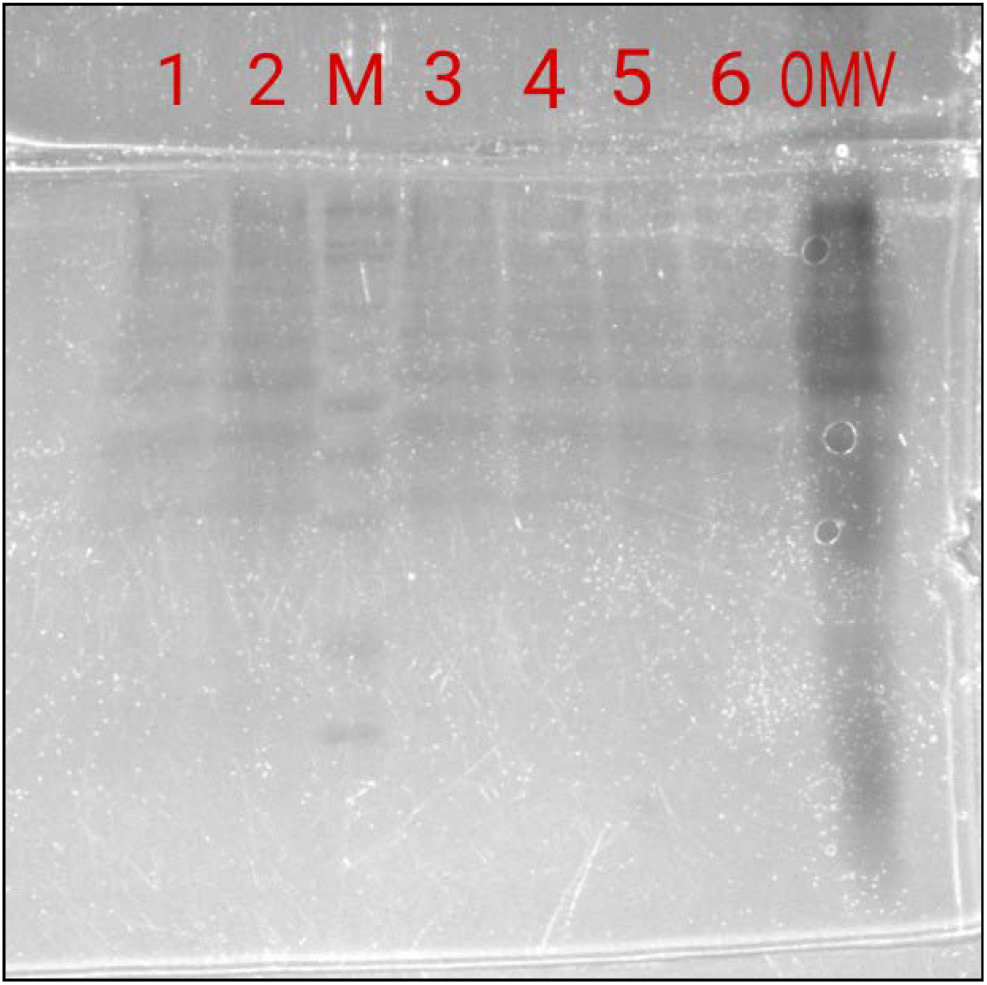
Protein profiles of vaccine formulations alongwith OMV of *Salmonella* Typhimurium by SDS-PAGE. Lane M: Hi Range Protein marker Lane 1: Chitosan nanoparticles-OMV conjugate * Lane 2: Chitosan nanoparticles-OMV conjugate ** Lane 3: PLG microparticles-OMV conjugate * Lane 4: PLG microparticles-OMV conjugate ** Lane 5: poly(anhydride) nanoparticles-OMV conjugate * Lane 6: poly(anhydride) nanoparticles-OMV conjugate ** Lane 7: Outer membrane vesicles * Predicted using RSM; ** control condition

Chitosan nanoparticle-OMV conjugation:

Entrapment efficiency % = 80.7391 + (3.5812× xA) - {0.5038×(xA)^2^}
PLG-OMV conjugation
Entrapment efficiency % = 98.26755+(0.81003× xA)-{0.09751×(xA)^2^}
Poly(anhydride) nanoparticle-OMV conjugation
Entrapment efficiency % = 99.38285+(0.92634 × x1) + (−0.38497 × x2) + (1.18236 ×
x1 × x2) -{ (0.20137 × (x1)^2^} - {(2.95451 × (x2)^2^}

Where xA= coded values of ratio, x1 and x2= coded values of pH and temperature.

### Revalidation of predicted entrapment efficiency in optimized conditions

The RSM predicted entrapment efficiencies of Ch-NP, PLG-MP and poly(anhydride) nanoparticles in the optimized conditions did not vary significantly from the one obtained experimentally in those conditions (table 3).

**Table 3:** Experimental revalidation of the RSM predicted optimum conditions.

| Nanoparticles/<br>Microparticles | RSM predicted<br>optimum condition | RSM predicted<br>entrapment<br>efficiency (%) | Mean<br>experimental<br>entrapment<br>efficiency (%)<br>(±SD) | t value |
| --- | --- | --- | --- | --- |
| Chitosan<br>nanoparticles-OMV conjugate | Ratio of<br>nanoparticles:OMV=<br>1:10 | 86.05 | 85.63±0.68 | -1.078<sup>NS</sup> |
| PLG<br>microparticles-OMV conjugate | Ratio of<br>microparticles:OMV=<br>1:9 | 99.94 | 99.88±0.08 | -1.309<sup>NS</sup> |
| poly(anhydride)<br>nanoparticles-OMV conjugate | pH=7,<br>Temp=24°C | 99.236 | 99.45±0.1 | 3.646<sup>NS</sup> |
| NS: p>0.05 |  |  |  |  |

### Effect of optimization on protein profile and immunopotential of nanoparticle/microparticle conjugates

The protein profile of the nanoparticles/microparticles—OMV conjugates prepared by RSM optimized conditions and the control conditions was not distinguishable in the SDS-PAGE. This indicates that the optimization of the conditions did not alter the protein profiles of the nanoparticles/microparticles—OMV conjugates.

There was no statistically significant difference between the immunogenic potential of nanoparticles/microparticles OMV conjugates prepared using RSM optimized conditions and the control conditions (table 4). Hence, it can be assumed that the optimized conditions are not detrimental to immunopotential of the conjugates. This may be due to the fact that the optimized conditions were within the physiological range of the proteins present in the OMVs.

**Table 4:** Anti-OMV IgM response of nanoparticles/ microparticles OMV conjugates in mice.

| Days | 7 |  | 14 |  | 7 |  | 14 |  | 7 |  | 14 |  |
| --- | --- | --- | --- | --- | --- | --- | --- | --- | --- | --- | --- | --- |
| Groups | Gr I | Gr II | Gr I | Gr II | Gr III | Gr IV | Gr III | Gr IV | Gr V | Gr VI | Gr V | Gr VI |
| Mean OD<sub>492</sub> | 0.24 | 0.19 | 0.37 | 0.32 | 0.52 | 0.46 | 0.49 | 0.43 | 0.67 | 0.62 | 0.43 | 0.53 |
| SE | 0.02 | 0.02 | 0.03 | 0.04 | 0.02 | 0.03 | 0.05 | 0.05 | 0.05 | 0.06 | 0.05 | 0.01 |
| Sig | NS |  | NS |  | NS |  | NS |  | NS |  | NS |  |

Gr I: poly(anhydride) nanoparticles-OMV conjugate\*
Gr II: poly(anhydride) nanoparticles-OMV conjugate\*\*
Gr III: PLG -OMV conjugate\*
Gr IV: PLG -OMV conjugate\*\*
Gr V: Chitosan nanoparticles-OMV conjugate\*
Gr VI: Chitosan nanoparticles-OMV conjugate\*\*
\* Control conditions, \*\* conditions predicted by RSM

## CONCLUSION

In the present investigation, our experimental findings revealed that the optimized conditions predicted by response surface method (RSM) having maximum OMV entrapment efficiency were pH and temperature of 7.0 and 24 °C respectively for poly(anhydride) nanoparticles and the ratio (NP:OMV) of 1:9 and 1:10 at 20-25°C and pH 7 for PLG microparticle and chitosan nanoparticle respectively. We believe that as the physical nature of OMVs of different Gram negative bacteria are similar, hence the optimized processes mentioned in this paper may work with OMVs of other bacteria also, however, further studies are required to confirm it.

## ACKNOWLEDGEMENT

This work was funded by DBT project entitled “Development of Nanoparticle/Microparticle Adjuvanted Subunit Oral Vaccine against Poultry Salmonellosis” and DPGS, Assam Agricultural University.

